# Physiological Condition Dependent Changes in Ciliary GPCR Localization in the Brain

**DOI:** 10.1101/2022.10.13.512090

**Authors:** Kathryn M. Brewer, Staci E. Engle, Ruchi Bansal, Katlyn K. Brewer, Kalene R. Jasso, Jeremy C. McIntyre, Christian Vaisse, Jeremy F. Reiter, Nicolas F. Berbari

## Abstract

Primary cilia are small immotile cellular appendages which mediate diverse types of singling and are found on most mammalian cell types including throughout the central nervous system. Cilia are known to localize certain G protein-coupled receptors (GPCRs) and are critical for mediating the signaling of these receptors. Several of these neuronal GPCRs have recognized roles in feeding behavior and energy homeostasis. Heterologous cell line and model systems like *C. elegans* and *Chlamydomonas* have implicated both dynamic GPCR cilia localization and cilia length and shape changes as key for signaling. However, it is unclear if mammalian ciliary GPCRs utilize similar mechanisms *in vivo* and under what physiological conditions these processes may occur. Here, we use the ciliary GPCRs, melanin concentrating hormone receptor 1 (MCHR1) and neuropeptide-Y receptor 2 (NPY2R) as model ciliary receptors to determine if dynamic localization to cilia occurs. We tested physiological conditions in which these GPCRs have been implicated such as feeding behavior, obesity, and circadian rhythm. Cilia were imaged using confocal microscopy and analyzed with a computer assisted approach allowing for unbiased and high throughput analysis of cilia. We analyzed GPCR positive cilia, cilia frequency as well as cilia length and receptor occupancy. Interestingly we observed changes in ciliary length, receptor occupancy, and cilia frequency under different conditions, but no consistent theme across GPCRs or brain nuclei was observed. A better understanding of the subcellular localization dynamics of ciliary GPCRs could reveal unrecognized molecular mechanisms regulating behaviors like feeding.

**Significance Statement:** Often, primary cilia localize specific G protein-coupled receptors (GPCRs) for subcellular signaling. Cell lines and model systems have indicated that cilia deploy dynamic GPCR localization and change their shape or length to modulate signaling. We used mice to assess neuronal cilia GPCRs under physiological conditions associated with both the receptors’ known functions and ciliopathy clinical features like obesity. We show that certain cilia with specific GPCRs appear to dynamically alter their length while others appear relatively stable under these conditions. These results implicate multiple themes across cilia GPCR mediated signaling and indicate that not all cilia modulate GPCR signaling using the same mechanisms. These data will be important for potential pharmacological approaches to target cilia GPCR-mediated signaling.

## Introduction

Cilia are near ubiquitous small microtubule-based cellular appendages critical for proper development and homeostasis where they coordinate specific signaling pathways (Reiter and Leroux 2017). Thus, cilia structure or function defects can result in a broad array of disorders with many clinical features (Reiter and Leroux 2017). These disorders, collectively known as ciliopathies, often are associated with neural developmental or behavioral deficits. For example, certain ciliopathies are associated with increased feeding behavior and obesity (Vaisse, Reiter et al. 2017, Engle, Bansal et al. 2021, Lee, Kang et al. 2022). Altered hypothalamic cilia signaling has been implicated in ciliopathies associated with obesity (Davenport, Watts et al. 2007, Loktev and Jackson 2013, Sun, Yang et al. 2021, Wang, Liu et al. 2021, Wang, Bernard et al. 2021).

Despite their clinical relevance and an understanding of cilia-mediated signaling in development, little is known about the roles of cilia on terminally differentiated neurons *in vivo* and how they influence mammalian behaviors. A growing and diverse set of G-protein coupled receptors (GPCRs) appear to preferentially localize to cilia, including specific GPCRs with known roles in feeding behavior and energy homeostasis, such as melanin concentrating hormone receptor 1 (MCHR1) and neuropeptide-Y receptor 2 (NPY2R) (Berbari, Johnson et al. 2008, Berbari, Lewis et al. 2008, Loktev and Jackson 2013).

During embryonic development, dynamic localization of signaling machinery and a GPCR (GPR161) to the ciliary compartment in a ligand-dependent manner is critical for proper hedgehog signaling (Mukhopadhyay, Wen et al. 2013, Hwang and Mukhopadhyay 2015, Pal, Hwang et al. 2016). In addition, *Chlamydomonas* and *C. elegans* utilize cilia length, shape, vesicular shedding, and receptor localization changes to mediate signaling (Mukhopadhyay, Lu et al. 2008, Olivier-Mason, Wojtyniak et al. 2013, Wang, Nikonorova et al. 2020, Wang, Nikonorova et al. 2021). Heterologous mammalian cell line data also clearly demonstrates the dynamic localization of ciliary GPCRs as a potential mechanism to mediate signaling, and ciliopathy mutations are associated with deficits in this process (Ye, Breslow et al. 2013, Nager, Goldstein et al. 2017, Phua, Chiba et al. 2017, Shinde, Nager et al. 2020).

In mammalian adult homeostasis, less is understood about how cilia mediate GPCR signaling in the central nervous system. The most well-studied examples are the photoreceptors and olfactory sensory neuron cilia, which mediate opsin/rhodopsin and odorant receptor signaling for vision and olfaction (Singla and Reiter 2006, Berbari, O’Connor et al. 2009). Here we sought to determine if cilia GPCR localization, frequency, and length dynamics change within brain regions associated with both the specific GPCRs function and ciliopathy-associated clinical features such as obesity. We focused on ciliary MCHR1 because it offers additional advantages of having a single receptor subtype in mice, is broadly expressed in the brain, and has been implicated in multiple physiological contexts such as feeding, metabolism, sleep, and reward (Pissios, Frank et al. 2008, Presse, Conductier et al. 2014, Blanco-Centurion, Luo et al. 2019, Dilsiz, Aklan et al. 2020).

## Materials and Methods

### Mice

All procedures were approved by the Institutional Animal Care and Use Committee at Indiana University-Purdue University Indianapolis. Adult C57Bl6/J mice were obtained from Jackson Laboratories, Bar Harbor, Maine (Stock #022409). Unless stated otherwise, mice were housed on a standard 12-hour light-dark cycle with *ad libitum* food and water.

### Feeding Conditions

<u>Fed</u> mice were allowed *ad libitum* access to food, <u>Fasted</u> mice had no food overnight (~16 hours), and <u>Refed</u> mice were given 4 hours of *ad libitum* access to food immediately after an overnight fast.

### Diet-induced Obesity

Mice were fed either a standard chow diet consisting of 13% fat, 58% carbohydrate, and 28.5% protein caloric content (LabDiet, #5001) or a calorie-rich, high-fat diet (HFD) consisting of 60% fat, 20% carbohydrate and 20% protein caloric content beginning at eight weeks of age (ResearchDiets, D12492). Mice were weighed weekly before proceeding to tissue analysis after 11 weeks on these diets and the onset of obesity.

### Circadian Time Point Conditions

Mice were randomly assigned to light or dark cycle perfusion groups. One hour before the light cycle (ZT23) and 4 hours before the dark cycle (ZT8), mice were anesthetized and perfused under their respective dark/light conditions.

### MCHR1 Antagonist Treatment

As previously described, mice were given an i.p. injection of 3 mg/kg MCHR1 antagonist, GW803430 (Tocris, #4242), or vehicle control for seven days, 3 hours after the start of the light cycle (Alhassen, Kobayashi et al. 2022). One week prior to the start of injections, mice were singly housed. Body weights were measured on the first day before injections to calculate the correct vehicle volume and dosage of GW treatment. MCHR1 antagonist was made fresh daily at a concentration of 0.5 mg/mL in 2 mL aliquots, (1 mg of GW, 8μL acetic acid,1.6 mL water, 125μL 2% Tween 80, 100μL 1N NaOH). Mice were weighed on the morning of the last treatment day (day seven) and perfused 60-90 minutes after the last injection.

### Fixation and Tissue Processing

Mice were anesthetized with 0.1 ml/ 10 g of body weight dose of 2.0% tribromoethanol (Sigma Aldrich) and transcardially perfused with PBS, followed by 4% paraformaldehyde in PBS (Electron Microscopy Sciences, #15710). Brains were postfixed in 4% paraformaldehyde for 4 hours at 4°C and then cryoprotected using 30% sucrose in PBS for 16–24 hours. Cryoprotected brains were embedded in Optimal Cutting Temperature compound (Fisher Healthcare, #4585) and sectioned at 15μm.

### Immunofluorescence

Sections were washed with PBS for 5 minutes, then permeabilized and blocked in a PBS solution containing 1% BSA, 0.3%TritonX-100, 2% (vol/vol) donkey serum, and 0.02% sodium azide for 30 minutes at room temperature. Sections were incubated with primary antibodies in blocking solution overnight at 4°C. Primary antibodies include anti-MCHR1 (rabbit pAB, Invitrogen, #711649, 1:250 dilution), anti-ACIII (chicken pAb, Encor, CPCA-ACIII, 1:1000 dilution), anti-mCherry (chicken pAb, Novus, NBP2-25158, 1:1000 dilution), anti-MCH (rabbit mAb, Abcam, #274415, 1:200 dilution). Sections were then washed with PBS before incubating with secondary antibodies for one hour at room temperature. Secondary antibodies include Donkey conjugated, Alexa Fluor 647, 488, (Invitrogen, 1:1000) against appropriate species according to the corresponding primary. All primary and secondary solutions were made in the blocking solution described above. Slides were then washed in PBS and stained with Hoechst nuclear stain (Invitrogen, #H3570) for 5 minutes at room temperature. Coverslips were mounted using SlowFade Diamond Antifade Mountant (Life Technologies, S36972).

### Mchr1 Antibody Validation

Brain sections from previously described Mchr1 knockout (*Mchr1^KO^*) and fluorescent reporter (*Mchr1^mCherry^*) mice were used for immunofluorescence to confirm the fidelity of the anti-MCHR1 antibody used throughout (**Supplemental Figure 1A** and **B**) (Jasso, Kamba et al. 2021).

### Confocal Imaging

All images were acquired using a Leica SP8 confocal microscope in resonant scanning mode using a 63x NA 1.4 objective. For all images collected, 16-bit image files were used for subsequent analysis.

### Image Analysis

Cilia analysis was performed as previously described (Bansal, Engle et al. 2021). Briefly, sum projection images from captured z-stacks were analyzed using the artificial intelligence (Ai) module, which had been trained to recognize cilia in brain section images. As part of the GA3 recipe, objects less than 1um in length were removed from the analysis. There were 4-5 mice per experimental condition, with four images captured per brain nuclei.

### Statistical Analysis

All statistical tests were performed using GraphPad Prism. All statistically significant observations are noted in the figures and specific tests used are named within the legends.

## Results

To understand if cilia GPCRs dynamically localize *in vivo* under physiological contexts associated with receptor activity, we initially chose to assess the known ciliary GPCR, MCHR1. We assessed its ciliary localization alone and in conjunction with the broadly expressed CNS ciliary membrane-associated adenylate cyclase 3 (ACIII) (Bishop, Berbari et al. 2007, Berbari, Johnson et al. 2008, Hsiao, Munoz-Estrada et al. 2021, Kobayashi, Tomoshige et al. 2021, Alhassen, Kobayashi et al. 2022). We confirmed our MCHR1 antibody immunofluorescence specificity by observing the loss of ciliary staining in a *Mchr1* knockout allele mouse brain and colocalization with a *Mchr1-mCherry* knock-in fusion allele mouse (**Supplemental Figure 1A** and **B**) (Jasso, Kamba et al. 2021). For our broader analysis of cilia localization, we utilized our recently reported computer-assisted approach for measuring cilia frequency, length, and fluorescence intensity (Bansal, Engle et al. 2021). This approach offers the advantages of being less biased and higher throughput.

As the melanin concentrating hormone (MCH) and MCHR1 signaling axis displays sexual dimorphism, our initial analysis compared cilia frequency, length, and fluorescence intensity in adult male and female mice (Messina, Boersma et al. 2006, Santollo and Eckel 2008). Surprisingly, we did not observe differences in these parameters in any of the brain regions assessed, including the hypothalamic arcuate (ARC) and paraventricular nucleus (PVN) and the nucleus accumbens (shell and core) (**Figure 1A** and **B**). As we did not observe differences between males and females, we continued the remaining studies utilizing adult males.

**Figure 1:**
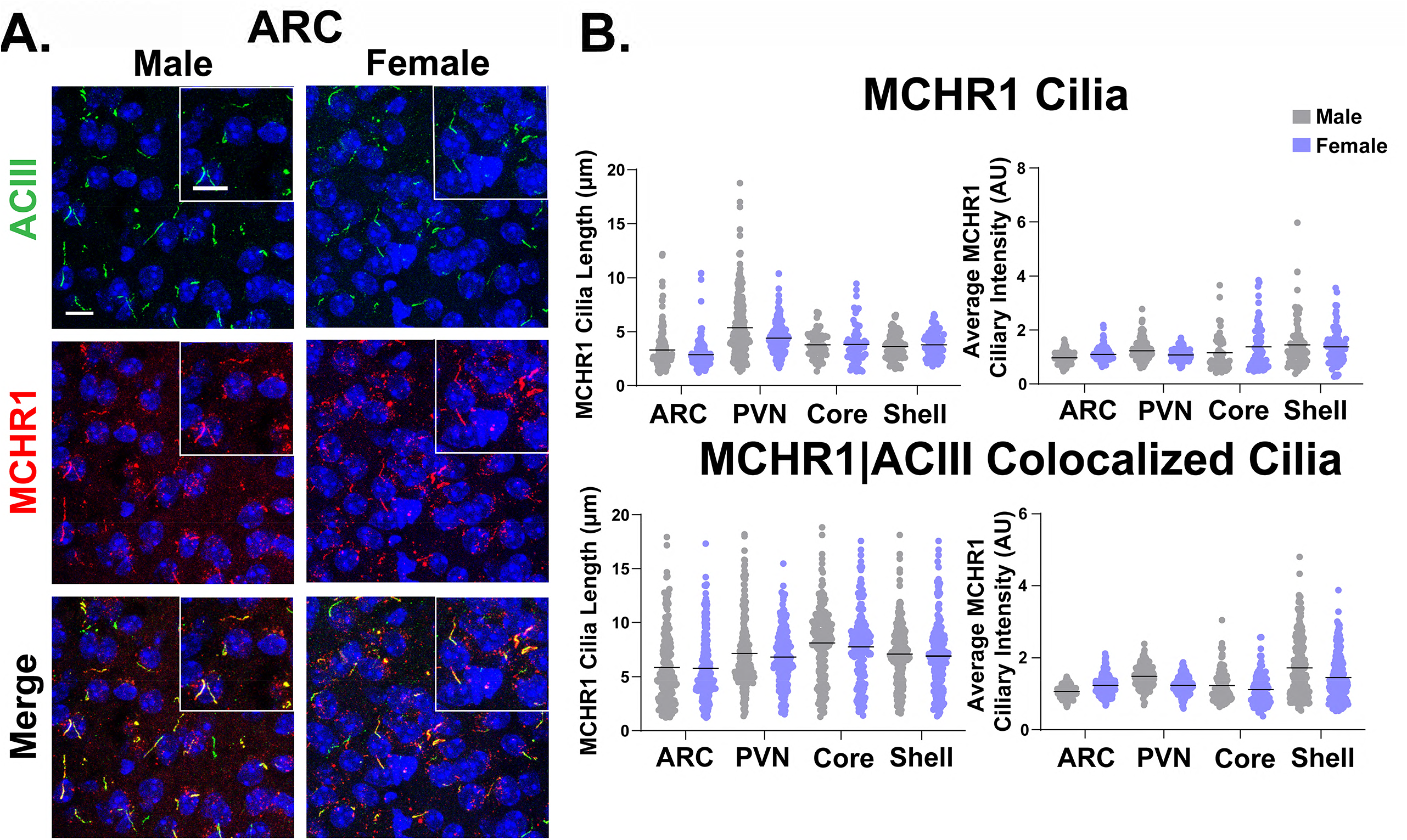
MCHR1 cilia localization is similar in adult male and female mice. **(A)** Representative immunofluorescence images of neuronal cilia (ACIII, green) and MCHR1 (red) in the arcuate nucleus (ARC) of males and females. Scale bars 10μm and Hoechst nuclei blue. **(B)** Mean MCHR1 cilia length and intensity in cilia with just MCHR1 (MCHR1 Cilia; top panels) or in cilia with both MCHR1 and ACIII (MCRH1|ACIII Colocalized Cilia; bottom panels) in the ARC, paraventricular nucleus (PVN) and the core and shell of the nucleus accumbens (nested t-test, p>0.05 for all male vs female comparisons in each region). N = 5 animals per group with an average of 250 cilia per brain nuclei of each animal analyzed.

MCHR1 function has been extensively implicated in feeding behaviors, body weight, and energy homeostasis (For a recent review, see (Al-Massadi, Dieguez et al. 2021)). Its ligand, MCH, is increased following acute fasting (Simon, Nemeth et al. 2018). Upon a 16-hour fast, we observed an increase in MCH ligand immunostaining in the lateral hypothalamus, the known site of MCH expression (**Figure 2A**) (Zamir, Skofitsch et al. 1986). We next assessed the impact of fasting on ciliary MCHR1 in both hypothalamic nuclei associated with this behavior, the ARC and PVN, and the nucleus accumbens, a site of high MCHR1 ciliary localization (Berbari, Lewis et al. 2008). Surprisingly, we only observed significant fasting-mediated changes in MCHR1 cilia length within the PVN (**Figure 2B** and **C**). To determine whether body weight and obesity can influence MCHR1 ciliary localization, we assessed the brains of high-fat diet induced obese mice (**Figure 3A**). Obesity did not influence cilia length or MCHR1 fluorescence intensity in the ARC, PVN, or accumbens (**Figure 3B** and **C**). In addition, neither fasting nor obesity altered MCHR1 cilia frequency observed in these brain regions (data not shown).

**Figure 2:**
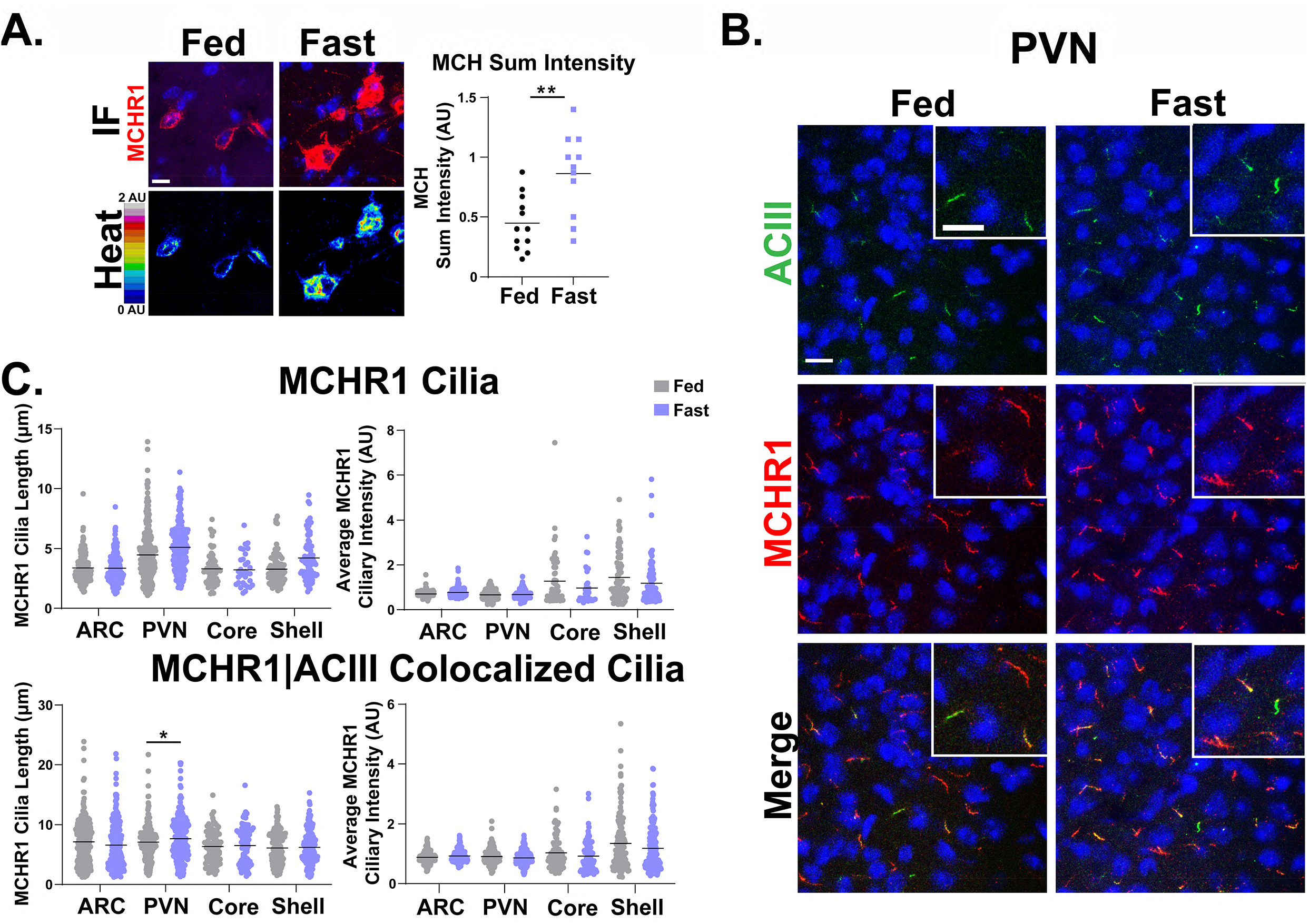
Acute feeding status alters MCHR1 length specifically in the PVN. **(A)** Melanin Concentrating Hormone (MCH) immunofluorescence staining (red) and intensity measurement (Heat) significantly increased under fasted conditions in the lateral hypothalamus, (student’s t-test, p=0.0024, 0.415AU±0.120). **(B)** Representative immunofluorescence images of neuronal cilia (ACIII, green) and MCHR1 (red) in the PVN of *ad libitum* fed (Fed) and fasted (Fast) animals. Scale bars 10μm and Hoechst nuclei blue. **(C)** Mean MCHR1 cilia length and intensity in cilia with just MCHR1 (MCHR1 Cilia; top panels) or in cilia with both MCHR1 and ACIII (MCRH1|ACIII Colocalized Cilia; bottom panels). Significant changes in MCHR1 cilia length were observed in the PVN (nested t-test, p=0.020, 0.62μm ±0.21). N = 5 animals per treatment group with an average of 200 cilia per brain nuclei of each analyzed. **\*** p<0.05, **\*\*** p<0.01.

**Figure 3:**
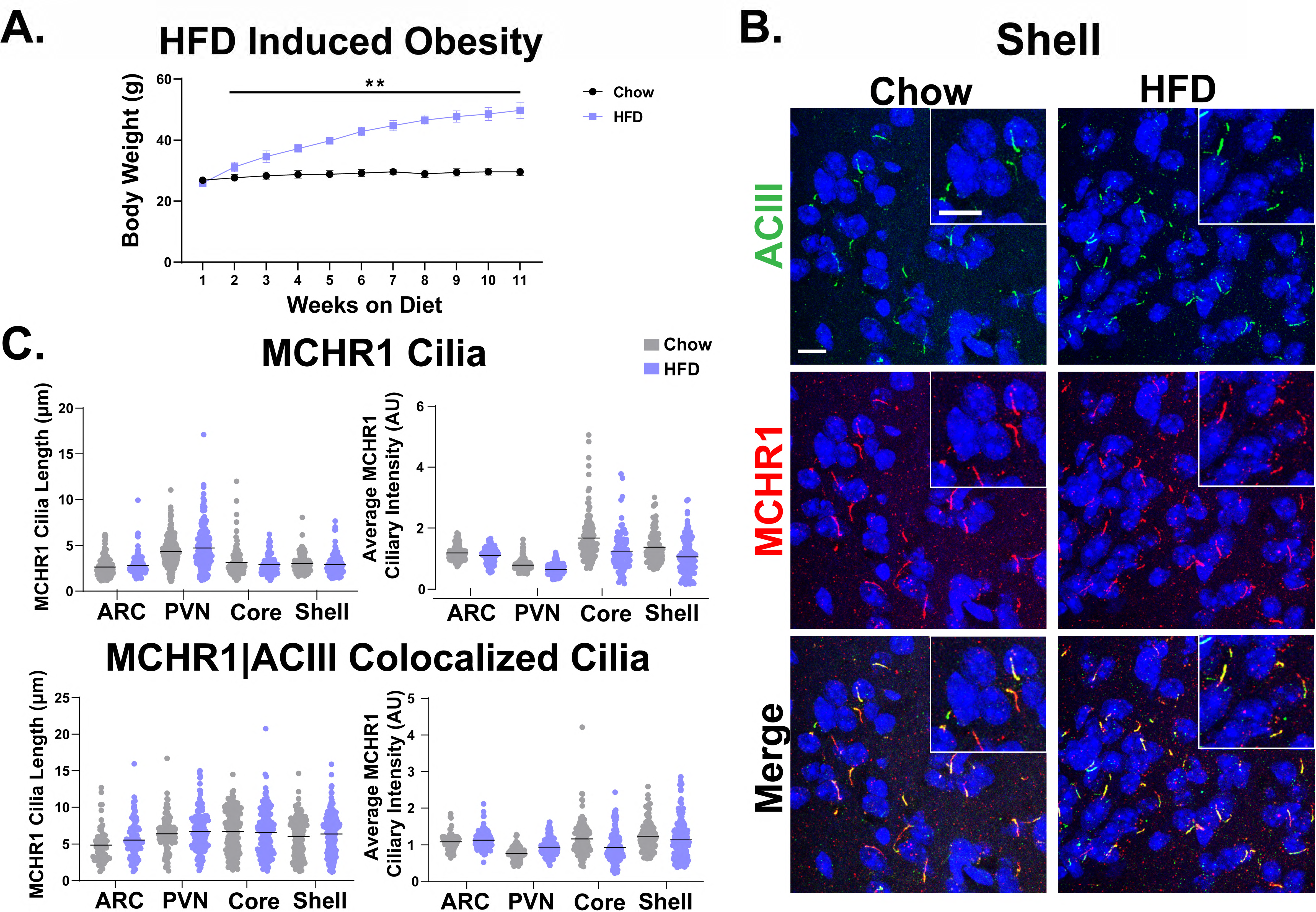
HFD induced obesity does not influence MCHR1 cilia localization. **(A)** High fat diet induced obese and chow fed controls animal body weights (student’s t-test p=0.008 at 2 weeks and is less than 0.0001 onward). **(B)** Representative immunofluorescence images of neuronal cilia (ACIII, green) and MCHR1 (red) in the ARC of control diet (Chow) and high fat diet (HFD) induced obese males. Scale bars 10μm and Hoechst nuclei blue. **(C)** Mean MCHR1 cilia length and intensity in cilia with just MCHR1 (MCHR1 Cilia; top panels) or in cilia with both MCHR1 and ACIII (MCRH1|ACIII Colocalized Cilia; bottom panels) (nested t-test, p>0.05). N = 5 animals per treatment group with an average of 250 cilia per animal and nuclei analyzed. ** p<0.01.

MCHR1 signaling has also been implicated in sleep/wake cycles (Blanco-Centurion, Luo et al. 2019). To determine if MCHR1 cilia localization changes with the light cycle, we assessed brains at ZT8 (light) and ZT23 (dark). We initially assessed the suprachiasmatic nucleus (SCN), the classic region involved in circadian rhythms and a where cilia length changes have recently cilia been implicated (Hastings, Maywood et al. 2018, Tu, Li et al. 2022). While we do not observe MCHR1 positive cilia in the SCN, we did note similar changes in ACIII cilia to those observed by Tu *et al* (**Figure 4A**). Interestingly, we also observed changes in MCHR1 cilia frequency in the ARC and PVN during the light/dark cycle (**Figure 4B**). In addition, MCHR1 cilia length in the shell of the accumbens appeared shorter in the dark cycle (ZT23) (**Figure 4C** and **D**). In the ARC the average MCHR1 fluorescence intensity was less in both MCHR1 cilia and those that had both MCHR1 and ACIII (**Figure 4D**).

**Figure 4:**
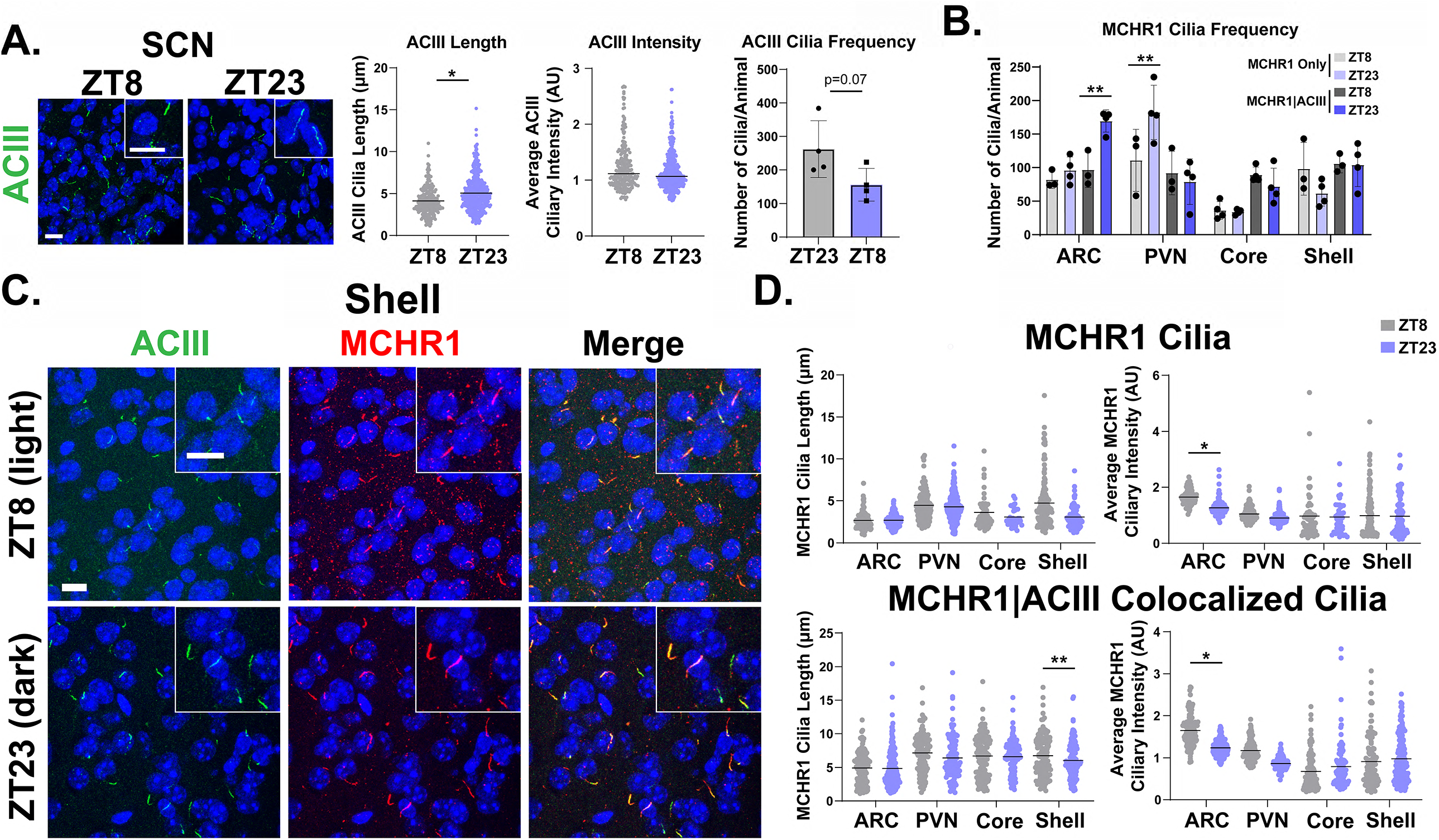
MCHR1 cilia localization is influenced by circadian rhythm. **(A)** Suprachiasmatic nucleus (SCN) ACIII cilia length at ZT23 (dark-cycle) (nested t-test, p=0.0291, 0.74μm±0.26). **(B)** MCHR1 cilia frequency in the ARC, PVN, Core and Shell at ZT8 and ZT23 for cilia with both MCHR1 and ACIII in the ARC and in cilia with only MCHR1 in the PVN at ZT23 (2-way ANOVA, ARC p=0.004, 73 cilia±20; PVN p=0.005, 70 cilia±20). **(C)** Representative immunofluorescence images of neuronal cilia (ACIII, green) and MCHR1 (red) in the shell at ZT8 (light cycle) and ZT23 (dark cycle) timepoints. Scale bars 10μm and Hoechst nuclei blue. **(D)** Mean MCHR1 cilia length and intensity in cilia with just MCHR1 (MCHR1 Cilia; top panels) or in cilia with both MCHR1 and ACIII (MCRH1|ACIII Colocalized Cilia; bottom panels. Significant decrease in MCHR1 cilia length in MCHR1|ACIII cilia in the shell and significant decreases in MCHR1 cilia flourescence intensity in the ARC at ZT23 (nested t-test, accumbens shell p=0.0089, −0.94 μm ±0.23; ARC p=0.0168, 0.386 AU±1.10; p=0.0147, −0.454 AU±1.24). N = 5 and 4 animals per treatment group, respectively, with an average of 200 cilia per animal and nuclei analyzed. * p<0.05, ** p<0.01.

After assessing multiple physiological conditions where MCHR1 function has been implicated, we next looked to see if overt pharmacological antagonism could influence MCHR1 ciliary localization. Injection of the antagonist, GW803430, for seven days resulted in significant changes in body weight (**Figure 5A**) (Alhassen, Kobayashi et al. 2022). MCHR1 antagonism increased the frequency of MCHR1 cilia in the ARC (**Figure 5B**). Antagonism also increased ciliary length in the accumbens core and shell for MCHR1 cilia and those that had both MCHR1 and ACIII (**Figure 5C** and **D**). Interestingly, in the ARC cilia length increases were observed in cilia with only MCRH1 (**Figure 5D**).

**Figure 5:**
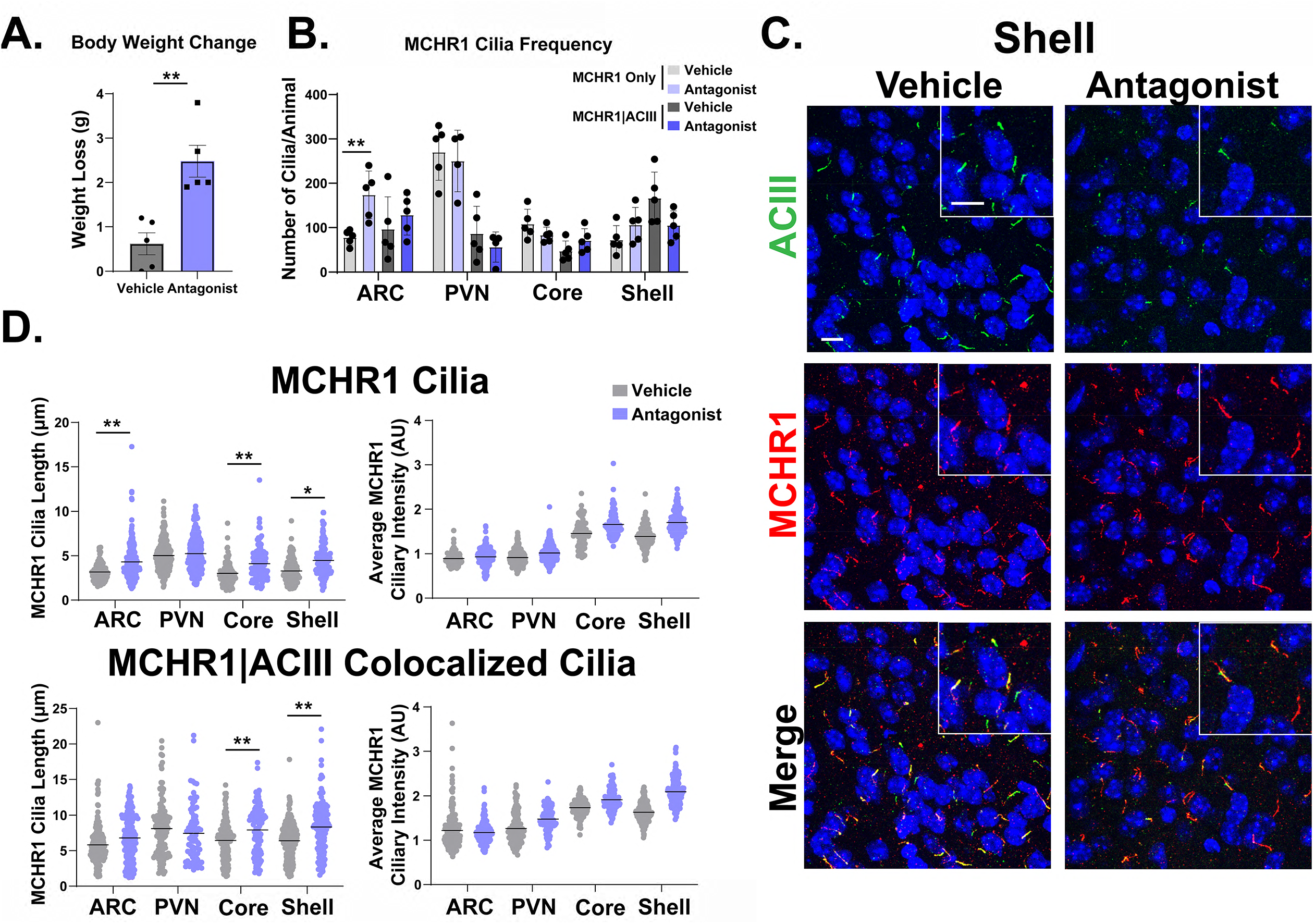
Antagonism alters MCHR1 length in the ARC and NA. **(A)** Antagonist treatment causes significant weight loss (student’s t-test, p=0.002, 1.860g ± 0.4368). **(B)** MCHR1 cilia frequency in the ARC, PVN, Core and Shell upon vehicle and antagonist treatment. Significant increase in MCHR1 only cilia upon antagonist treatment in the ARC (2-way ANOVA, p=0.008, 96 cilia±28). **(C)** Representative immunofluorescence images of neuronal cilia (ACIII, green) and MCHR1 (red) in the shell of control (Vehicle) and MCHR1 antagonist (Antagonist) treated animals. Scale bars 10μm and Hoechst nuclei blue. **(D)** Mean MCHR1 cilia length and fluorescence intensity in cilia with just MCHR1 (MCHR1 Cilia; top panels) or in cilia with both MCHR1 and ACIII (MCRH1|ACIII Colocalized Cilia; bottom panels) Significant changes in cilia length for both MCHR1 and MCHR1|ACIII cilia in the ARC, core, and (MCHR1 cilia nested t-test, ARC p=0.0089, 0.94μm±0.23; accumbens core p=0.0033, 0.97μm±0.31; accumbens shell p=0.0224, 0.89μm±0.31. MCRH1|ACIII Colocalized Cilia nested t-test, accumbens core p=0.0003, 1.47μm±0.24; accumbens shell p<0.0001, 1.70μm±0.22 respectively). N = 5 animals per treatment group with an average of 250 cilia per animal and nuclei analyzed. * p<0.05, ** p<0.01.

To determine if these results are specific to MCHR1 or perhaps applicable to multiple neuronal ciliary GPCRs, we assessed the localization of neuropeptide-Y receptor 2 (NPY2R), another GPCR known to localize to cilia in the ARC and be involved in feeding behaviors (Loktev and Jackson 2013). Acute fasting increases the levels of the NPY2R ligand, neuropeptide-Y (NPY) (Yasrebi, Hsieh et al. 2016). Thus, we sought to assess both MCHR1 and NPY2R upon fasting and refed states. We only observed significant changes in MCHR1 cilia length between the fasted and refed conditions (**Figure 6A** and **B**). However, we observed significant cilia length changes in NPY2R cilia and those with both NPY2R and ACIII between *ad libitum* fed, fasting, and refed conditions (**Figure 6C** and **D**).

**Figure 6:**
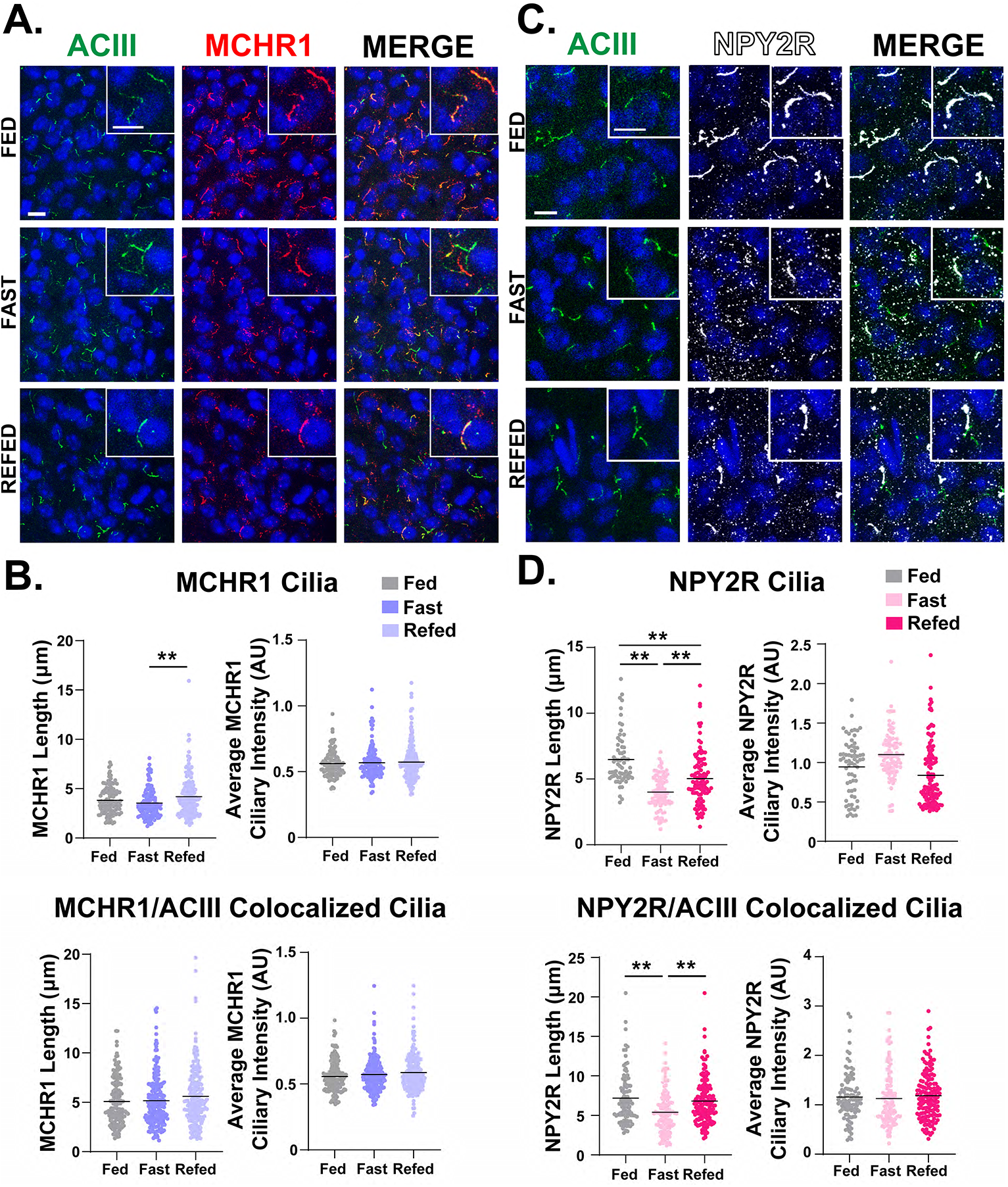
NPY2R cilia length and localization changed under different feeding statuses in the ARC. **(A)** Representative immunofluorescence images of neuronal cilia (ACIII, green) and MCHR1 (red) in the ARC of *ad libitum* fed (Fed), overnight fasted (Fast) and 4 hours post refeeding after fast (Refed) conditions. **(B)** Mean MCHR1 cilia length and intensity in cilia with just MCHR1 (MCHR1 Cilia; top panels) or in cilia with both MCHR1 and ACIII (MCRH1|ACIII Colocalized Cilia; bottom panels) in Fed, Fast and Refed animals. Significant increase in MCHR1 cilia length upon refeeding (MCHR1 nested 1-way anova, p=0.004, 0.654μm±0.201). **(C)** Representative immunofluorescence images of neuronal cilia (ACIII, green) and NPY2R (white) in the ARC of Fed, Fast and Refed conditions. **(D)** Mean NPY2R cilia length and intensity in cilia with just NPY2R (NPY2R Cilia; top panels) or in cilia with both NPY2R and ACIII (NPY2R|ACIII Colocalized Cilia; bottom panels) in Fed, Fast and Refed animals. NPY2R cilia in the ARC significantly change length upon Fed, Fast, and Refed conditions (nested 1-way ANOVA, p<0.001, −2.47μm±0.29; p<0.0001,1.45 μm ±0.28; p=0.0002, −1.02 μm ±0.25; respectively). Cilia in the ARC that are positive for both NPY2R and ACIII (NPY2R|ACIII Colocalized Cilia) are also significantly shorter upon fasting and remain slightly shorter in the Refed condition compared to Fed (nested t-test, p<0.0001, −1.80μm±0.41; p=0.0003, 1.43μm±0.37, respectively). Each data point represents a cilium. For all images, scale bars 10μm and Hoechst nuclei blue. N = 5 animals per group with an average of 250 cilia per animal. * p<0.05, ** p<0.01

## Discussion

Cilia are recognized as mediators of diverse signaling pathways, yet many questions remain unanswered regarding how they coordinate signaling. In cell line and heterologous expression systems *in vitro*, dynamic localization of receptors to the cilia membrane has been reported for a number of ciliary GPCRs, including MCHR1 (Ye, Nager et al. 2018). *In vivo* dynamic localization to the cilia as a means of signaling control has been best described for the cilia mediated hedgehog signaling during development (Bangs and Anderson 2017). We sought to determine if cilia broadly deploy dynamic GPCR localization *in vivo* to mediate signaling. We chose a ciliary receptor associated with several physiological states and phenotypes, including sexual dimorphic expression, acute feeding behavior, energy homeostasis and sleep (Al-Massadi, Dieguez et al. 2021). Our initial assessment of MCHR1 focused on the hypothalamus for a number of reasons. Ciliopathies are known to have deficits in hypothalamic control of energy homeostasis (Davenport, Watts et al. 2007, Sun, Yang et al. 2021, Wang, Bernard et al. 2021). MCHR1 fails to localize properly in obese ciliopathy models of Bardet-Biedl syndrome (BBS) (Berbari, Lewis et al. 2008) and most importantly *Mchr1* expression is observed in several hypothalamic nuclei under baseline conditions (Engle, Antonellis et al. 2018). In addition, MCHR1 signaling has been extensively implicated in feeding behavior, energy homeostasis, and metabolism. Agonism or activation of the pathway is associated with increases in food intake, and loss-of-function alleles or pharmacological antagonism associated with weight loss (for a recent review on MCH and MCHR1 signaling see: (Al-Massadi, Dieguez et al. 2021)).

We chose an antibody staining approach combined with a computer assisted analysis as this combination was the best way to detect endogenous ciliary MCHR1 in an unbiased and high throughput manner allowing us to observe hundreds of cilia per animal (Bansal, Engle et al. 2021, Jasso, Kamba et al. 2021).

We were surprised to find that our analysis revealed that MCHR1 ciliary localization remained largely fixed across males and females, upon fasting and diet induced obesity, with only subtle significant changes observed in cilia length. Our observation that ACIII cilia length changes within the SCN depending on the light or dark cycle as recently reported in a pre-print, assured us that our analysis could detect broad scale changes in cilia lengths, frequency and localization (Tu, Li et al. 2022). It was interesting that we also detected length changes in cilia with both MCHR1 and ACIII in the shell of the nucleus accumbens at different circadian time points (Becker-Krail, Walker et al. 2022). This suggests the potential for cilia mediated signaling changes broadly in the brain based on light conditions.

Pharmacological MCHR1 antagonism demonstrated the most substantial changes in both cilia length and intensity across different brain regions but this approach may not be physiologically relevant. However, this result is in line with what cilia have been proposed to do when their GPCR associated signaling system is saturated or overwhelmed by changing their lengths and shedding cilia specific vesicles (Nager, Goldstein et al. 2017, Phua, Chiba et al. 2017). These phenomena have been directly observed for cilia in BBS cell models (Nager, Goldstein et al. 2017). It remains to be seen how common cilia length regulation and vesicular shedding is deployed as a means of cilia mediated signaling *in vivo*. It is possible that both are important process, but that under normal physiological conditions they remain challenging to detect in mammalian systems *in vivo* with currently available tools.

To further explore the possibility that other cilia GPCRs could be relatively stationary *in vivo*, we investigated another hypothalamic ciliary GPCR under physiological conditions in which it has been implicated; NPY2R and feeding status (Loktev and Jackson 2013). Interestingly, for NPY2R, we observed significant changes in cilia length for cilia with just NPY2R in fasted and refed conditions compared to *ad libitum* fed animals. These data suggest that NPY2R cilia are more dynamic upon acute changes in feeding when compared to MCHR1 cilia. Overall, these data further point to the potential that many ciliary GPCRs may need to be assessed independently for how their signaling is mediated *in vivo*.

Together our results demonstrate that dynamic localization to the ciliary compartment may not apply to some physiological conditions *in vivo* or be a common theme across ciliary GPCRs. Our results also suggest that only specific ciliary GPCRs utilize length control as a mechanism to mediate signaling, as may be the case for NPY2R but not MCHR1. Finally, our results also demonstrate that localization across different brain regions and nuclei that all possess the same ciliary GPCR do not show the same length regulation dynamics. For example, even upon supraphysiological antagonism of MCHR1, we did not observe the same changes in cilia length and localization in all brain regions analyzed. Ultimately, a comprehensive understanding of how cilia mediate GPCR signaling could provide therapeutic opportunities for cilia-receptor ligands in conditions like obesity.

## 13. Acknowledgements

Authors would like to thank Lata Balakrishnan for critical review

---

**Supplemental Figure 1:**
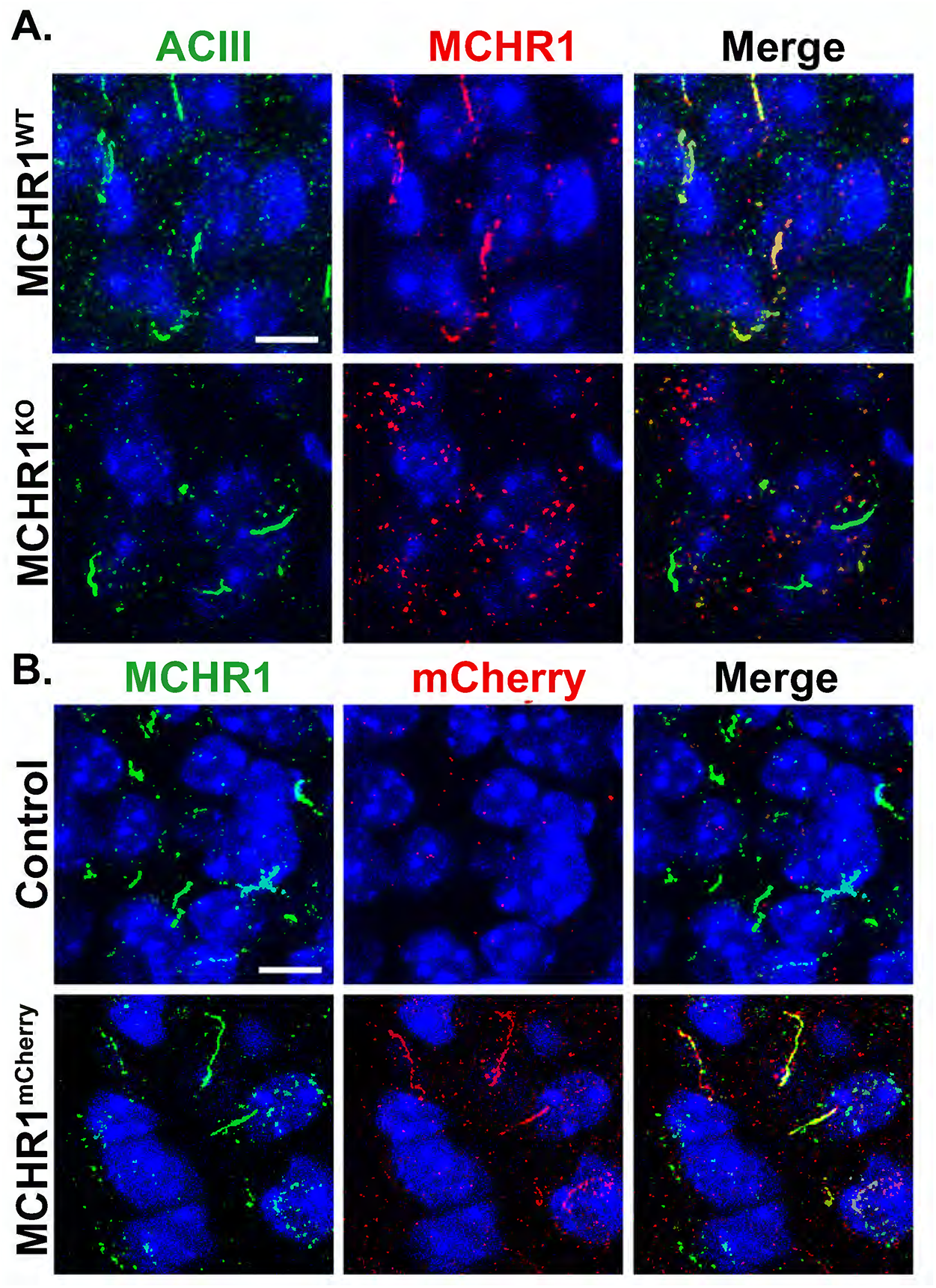
Antibody validation in MCHR1^KO^ and Mchr1^mCherry^ fusion allele animals. **(A)** MCHR1 knockout mice show ACIII positive cilia but show no MCHR1 positive cilia. **(B)** MCHR1 mCherry tagged mice show colocalization of MCHR1 and mCherry tag positive cilia. Scale bars 10μm and Hoechst nuclei blue. N = 3 animals per genotype.

